# Aging selectively dampens oscillation of lipid abundance in white and brown adipose tissue

**DOI:** 10.1101/2020.09.04.283226

**Authors:** Ntsiki M. Held, M. Renate Buijink, Hyung L. Elfrink, Sander Kooijman, Georges E. Janssens, Frédéric M. Vaz, Stephan Michel, Riekelt H. Houtkooper, Michel van Weeghel

**Author notes:** **Corresponding authors:** Riekelt Houtkooper; Michel van Weeghel; Laboratory Genetic Metabolic Diseases, Amsterdam UMC-AMC, 1105 AZ Amsterdam, The Netherlands.

## Abstract

Lipid metabolism is under the control of the circadian system and circadian dysregulation has been linked to obesity and dyslipidemia. These factors and outcomes have also been associated to, or affected by, the process of aging. Here, we investigated whether murine white (WAT) and brown (BAT) adipose tissue lipids exhibit rhythmicity and if this is affected by aging. To this end, we have measured the 24 hour lipid profiles of WAT and BAT using a global lipidomics analysis of >1100 lipids. We observed rhythmicity in nearly all lipid classes including glycerolipids, glycerophospholipids, sterol lipids and sphingolipids. Overall, ~22% of the analyzed lipids were considered rhythmic in WAT and BAT. Despite a general accumulation of lipids upon aging the fraction of oscillating lipids decreased in both tissues to 14% and 18%, respectively. Diurnal profiles of lipids in BAT appeared to depend on the lipid acyl chain length and this specific regulation was lost in aged mice. Our study revealed how aging affects the rhythmicity of lipid metabolism and could contribute to the quest for targets that improve diurnal lipid homeostasis to maintain cardiometabolic health during aging.

## 1. Introduction

Circadian clocks are responsible for the daily regulation of many physiological functions such as the sleep-wake cycles, energy expenditure, food intake and hormone secretion. The central clock of mammals is located in the suprachiasmatic nucleus (SCN) of the hypothalamus and is entrained by the light-dark cycle. It synchronizes the many clocks in the periphery. In return, the peripheral clocks, for instance those of the muscle, liver, gut and adipose tissue provide feedback to the central clock, for instance through release of signaling molecules, metabolite levels and the cellular redox state [1]. At the molecular level, the circadian clocks consist of transcription factors that create a transcription-translation feedback loop. The core clock transcription factors BMAL1 and CLOCK create the positive, and the PER and CRY proteins the negative feedback loop that together drive rhythmic gene expression and metabolism [2]. Circadian rhythms and metabolism are closely related as oscillating metabolic redox processes and intermediary metabolites provide feedback cues that synchronize peripheral clocks [3]. Misalignment of these components of the circadian system have been associated with age-related cardiometabolic diseases such as insulin resistance and obesity [4]. Circadian rhythms are also involved in the process of aging. Previously, BMAL1 deficiency has been shown to reduce the lifespan of mice, and dysregulated metabolism due to circadian misalignment is believed to accelerate aging [5,6]. On the other hand, several parts of the circadian system are known to decline in function during aging [7]. This is especially true for the light entrainment of the central clock, but also the sleep-wake cycles and certain hormonal feedback cues from the peripheral clocks change [8]. There is limited research on the effect of aging on peripheral clocks with particular focus on how this affects lipid metabolism.

Adipose tissue is a metabolic organ under strict circadian control, which was found to be sensitive to ageing. For instance, the PER2 protein of the circadian transcription machinery controls the expression of PPARγ, a master regulator of adipose function [9]. This suggests that many enzymes involved in adipocyte differentiation, lipid metabolism, glucose homeostasis and also adipokine secretion are regulated in a circadian manner [10]. Indeed, PER2 ablation affects the hydrolysis of triglycerides, triglyceride synthesis and increases fatty acid oxidation [9]. There are two main types of adipose tissue, i.e. white adipose tissue (WAT) and brown adipose tissue (BAT), which have distinct functions and are differentially affected by the circadian system and aging [11]. WAT stores energy in the form of lipids as triglycerides but also releases important adipokines such as leptin and adiponectin. On the contrary, BAT uses its stored lipids to dissipate energy as heat during non-shivering thermogenesis [12]. In general, adiposity increases upon aging. This is mostly reflected by an increase in WAT content. However, lipodystrophy is also a common feature of aging as BAT volume and activity decreases, thereby contributing to the age-related incidence of insulin resistance [13].

Previous studies in WAT, BAT and liver have shown that aging does not alter expression of the core clock components. Rather, aging is associated to altered rhythmic expression of downstream genes [14,15]. Many of these rhythmic genes are involved in metabolism, and therefore rhythmicity of metabolites might be a better readout to explore the effect of aging on circadian oscillations. Previously, we have shown that lipid metabolism in human muscle tissue exhibits day-night rhythmicity [16]. In the present study, we have compared the rhythmicity of the lipidome of WAT vs BAT. Furthermore, we compared the lipidome of WAT and BAT of young and old mice. We found that lipids tended to accumulate upon age, while the number of oscillating lipids reduced with age. Not only did the number of oscillating lipids decrease, we also found changes in the rhythmic profile, i.e. time of day that lipids peak.

## 2. Materials and Methods

### 2.1 Animals and housing

A total of 24 young (about 2 months) and 24 old (about 24 months) C57BL/6 male mice were acclimated to a set 12:12-h light/dark cycle. Animal handling was performed at the Leiden University Medical Center (LUMC), Leiden, the Netherlands and was previously approved by the institutional ethics committee on animal care and experimentation conforming to Dutch law (DEC 12250). The mice were group housed with chow (Special Diets Services, Essex, UK) and water available ad libitum. Zeitgebers time (ZT) 0 corresponded to the time lights were turned on (50-100 lx; Osram true light TL) and ZT12 to the time lights were turned off in the animal facility. On the day of the experiment, mice were killed by decapitation at 4-hr intervals around the clock, i.e. ZT2, 6, 10, 14, 18 and 22. Interscapular brown adipose tissue (BAT) and epididymal white adipose tissue (WAT) were removed and weighed prior to freezing in liquid nitrogen and were stored at −80°C until further use.

### 2.2 Lipid extraction

The adipose tissues were homogenized in ultra-pure ice-cold water using the TissueLyser system (Qiagen). Subsequently, the homogenates were sonicated in an ice-cold water bath for 10 min. Protein concentrations of the homogenates were determined using the BCA protein assay kit (Pierce). Lipids were extracted using a single-phase extraction, as described previously [17]. In brief, 1.5 mL of chloroform/methanol (1:1, v/v) and an internal standards mixture specifically adjusted to WAT or BAT was added to the tissue homogenate (the equivalent of 1 mg of protein). All lipid internal standards used are listed in **Table 1** internal standards (all from Avanti Polar Lipids, Alabaster, AL, USA).

**Table 1.**
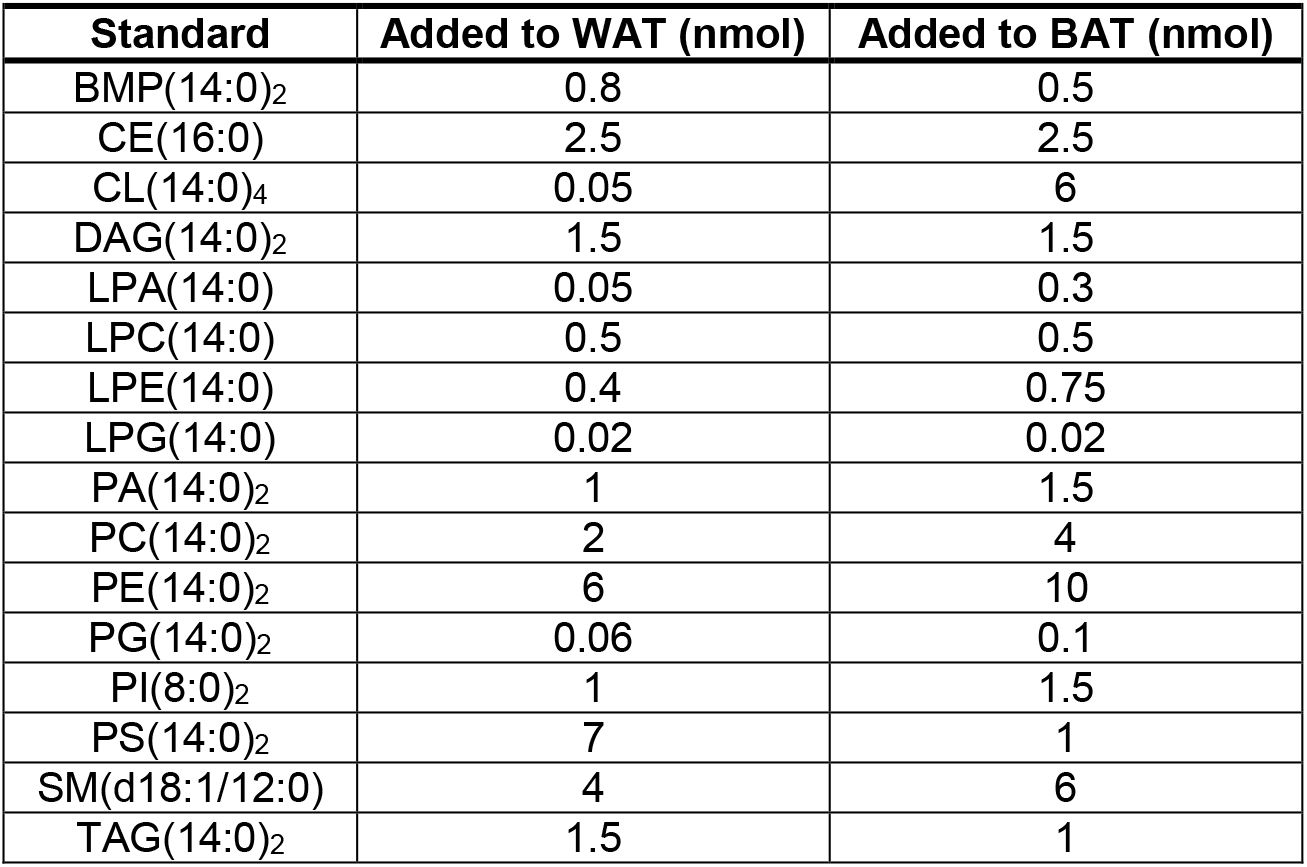
Added internal standards.

Subsequently, the mixture was sonicated in a water bath for 5 min, followed by centrifugation at 4 °C (16000x*g* for 5 min). The liquid phase was transferred to a glass vial and evaporated under a stream of nitrogen at 60 °C. Subsequently, the residue was dissolved in 150 μL of chloroform/methanol (9:1, v/v).

### 2.3 Ultra Performance Liquid Chromatography-High Resolution Mass Spectrometry (UPLC-HRMS)

Lipidomics analysis was performed as described previously with minor changes described in this section [18]. The UPLC system consisted of an Ultimate 3000 binary HPLC pump, a vacuum degasser, a column temperature controller, and an autosampler (Thermo Scientific, Waltham, MA, USA). For normal phase separation of lipids, 2 μL lipid extract was injected on a LiChroCART 250–4 LiChrospher^®^ Si 60 (5 μm) (Merck) maintained at 25 °C. For lipid separation a linear gradient consisting of solvent A (methanol/water, 85:15, v/v) and solution B (chloroform/methanol, 97:3, v/v) was used. Solvent A and B contained 5 and 0.2 mL of 25% (*v*/v) aqueous ammonia per liter of eluent, respectively. The gradient (0.3mL/min) was as follows: t = 0–1 min: 10%A; t= 1–4min: 10%A-20%A; t = 4–12min: 20%A-85%A; t=12–12.1 min: 85%A - 100%A; t= 12.1–14.0min: 100%A; t= 14–14.1 min: 100%A-10%A and t= 14.1–15 min: 10%A. For reverse phase separation of lipids, 5 μL lipid extract was injected onto an ACQUITY UPLC HSS T3 (1.8 μm, Waters) maintained at 60 °C. For lipid separation a linear gradient consisting of solvent A (methanol/water, 40:60, v/v) and solution B (methanol/isopropanol, 10:90, v/v) was used. Solutions A and B both contained 0.1% formic acid and 10 mM ammonia. The gradient (0.4 mL/min) was as follows: t = 0–1 min: 100%A; t= 1–16 min: 80%A; t= 16–20 min: 0%A; t = 20–20.1 min: 0%A; t = 20.1–21.0 min: 100%A. A Q Exactive Plus Orbitrap mass spectrometer (thermo Scientific) was used in the negative and positive electrospray ionization mode using nitrogen as nebulizing gas. The spray voltage was 2500 V, and the capillary temperature was 256 °C. S-lens RF level: 50, Auxiliary gas: 11, Auxiliary gas temperature 300°C, Sheath gas: 48 au, Sweep cone gas: 2 au. Mass spectra of lipids were acquired in both scan modes by continuous scanning from m/z 150 to m/z 2000 with a resolution of 280,000 full width at half maximum (FWHM) and processed using an in-house developed lipidomics pipeline written in the R programming language (http://www.r-project.org). The identified peaks were normalized to the intensity of the internal standard for each lipid class.

### 2.4 Statistical analysis

Significance levels of differences between groups were evaluated using either an unpaired two-tailed Student’s t-test or one-way analysis of variance (ANOVA) with Bonferroni post-hoc test with *P*-value <0.05. For the determination of rhythmicity in lipid concentration, only the lipids with relative abundance of >0.05 averaged over all 6 time points were taken into account. Rhythmicity of lipid levels were analyzed using the nonparametric test JTK_CYCLE, incorporating a window of 10-28 hr [19]. Lipid levels were considered rhythmic with a Bonferroni corrected *P*-value of <0.05. Subsequent statistical analyses and plotting were done in a R environment using the ggplot2, ropls and mixOmics packages [20–22]. The Venn diagram was created using the web-based tool Venny [23].

## 3. Results

### 3.1 Lipid profiling in WAT and BAT of young and aged mice

To establish the effects of aging on periodic regulation of lipid metabolism we characterized epididymal WAT and interscapular BAT of 2 months (‘young’) and 24 months (‘old’) old mice that were sacrificed in 24 hours at 4hrs intervals. WAT and BAT were harvested and characterized using UPLC-HRMS-based lipidomics. Data were processed by an in-house developed lipidomics pipeline and the day-night rhythmicity of lipids was analyzed with JTK_CYCLE (**Figure 1A**). We detected lipids belonging to 19 lipid classes, among these were sterol lipids (CE); glycerolipids (DAG, TAG and TG[O]); glycerophospholipids (BMP, CL, MLCL, PC, PC[O], LPC, PE, PE[O], LPE, LPE[O], PG, LPG and PI) and sphingolipids (SMd and SMt) (**Figure 1B**). In total, we found 1084, 1152, 1258 and 1291 distinct lipids in WAT young, BAT young, WAT old and BAT old mice, respectively (**Figure 1C-F**). The overall lipid class distribution was similar between groups. Although our method is not quantitative with respect to interclass comparisons (e.g. TAG vs PC), the relative abundance of the same lipid class across tissues can be compared (TAG in WAT vs TAG in BAT). The main difference between WAT and BAT was observed in cardiolipin (CL) content, a specific glycerophospholipid class that is highly enriched in the mitochondrial membrane where it plays an important role in the structural organization [24]. Given that the mitochondrial content in BAT exceeds that of WAT [25] it is not surprising that the CL content was twice as high in BAT compared to WAT (**Figure 1C,E** vs **D,F**). CL content is not only important for mitochondrial function, but also for BAT thermogenic capacity, which is known to decline with age [13,26–28]. Therefore, we next set out to determine the effect of aging on the individual lipid species.

**Figure 1:**
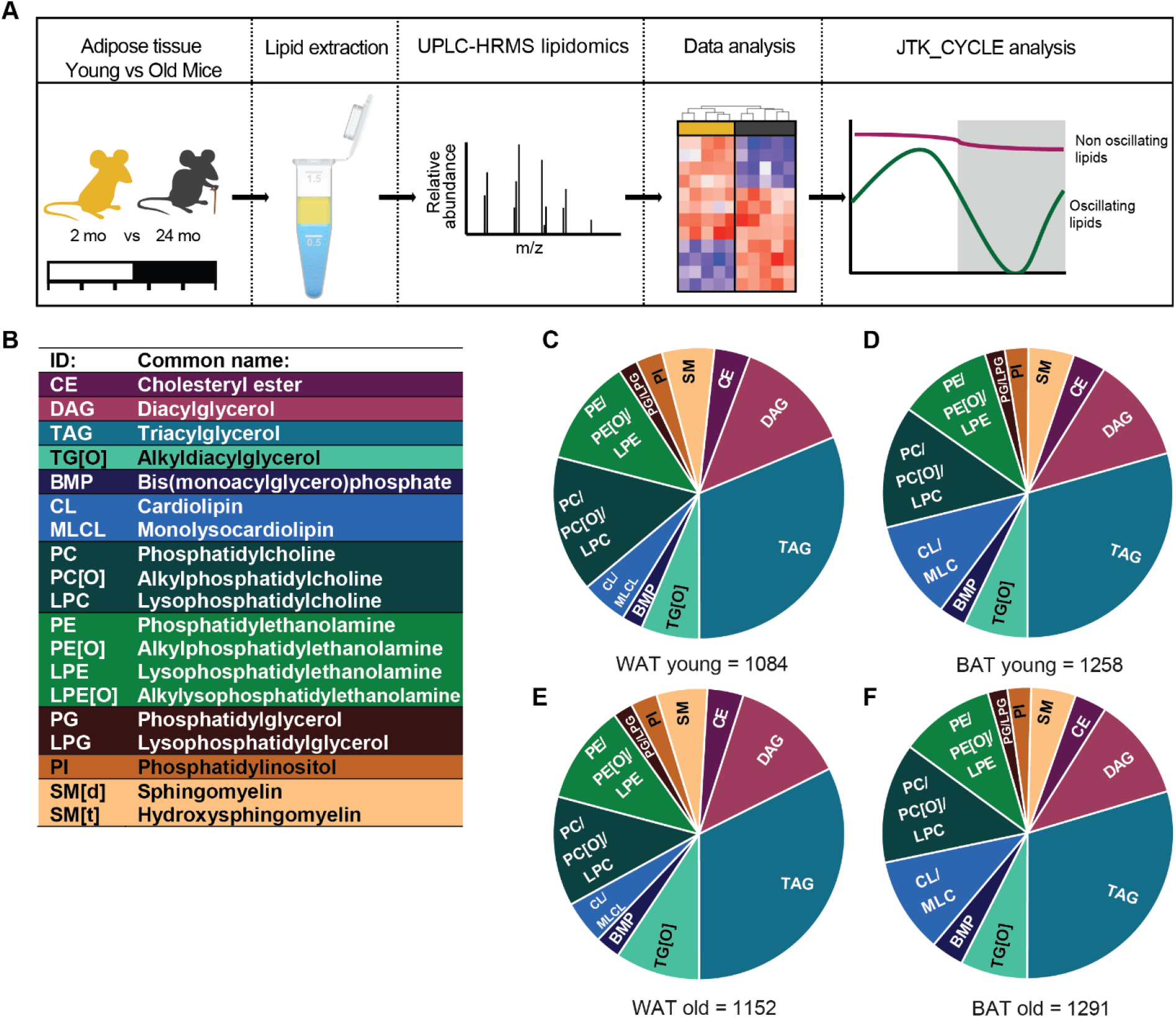
Lipid profiling in WAT and BAT of young and aging mice. (**A**) Experimental design: 2 months and 24 months old mice were sacrificed every 4 hours; WAT and BAT was harvested and lipids were extracted; lipidomics and data analysis was performed; and the obtained lipid profiles were analyzed with JTK_CYCLE analysis for detection of oscillating lipids. (**B**) The abbreviations and common names of the 19 detected lipid classes. The lipid composition of young mice in WAT (**C**), BAT (**D**), and of aged mice in WAT (**E**) and in BAT (**F**).

### 3.2 Aging induces an overall accumulation of lipids in WAT and specific lipid accumulations in BAT

First, we selected one time point to compare the lipid profiles of young and aged mice in WAT and BAT. The time point that we chose was ZT14 (early night). In this period mice are more physically active and also adipose tissue is more metabolically active [29]. Despite the similarities in overall lipid class distribution, the WAT lipidome of young mice is clearly different when compared to that of aged mice (**Figure 2A**). Most lipids classes were increased in old mice (**Figure 2 B-E**). Alkyldiacylglycerol, TG[O], was almost absent in young mice and increased 16-fold in aged mice. The same is true for DAGs, which increased by 6-fold upon aging (**Figure 2C**). Interestingly, TAGs were not significantly increased, although TAGs composed of long (>C22) monounsaturated or diunsaturated fatty acids showed a trend of higher levels in aged mice (**Figure 2B**). In the group of glycerophospholipids, the four-fold increase in BMPs and the two-fold increase in SM[t] was most striking. Also, alkylated lipids including PC[O], PE[O] and LPE[O] were all significantly increased ~2-fold. LPE appeared to be the only lipid class that was lower in aged mice, although this was not statistically significant (**Figure 2D**).

**Figure 2:**
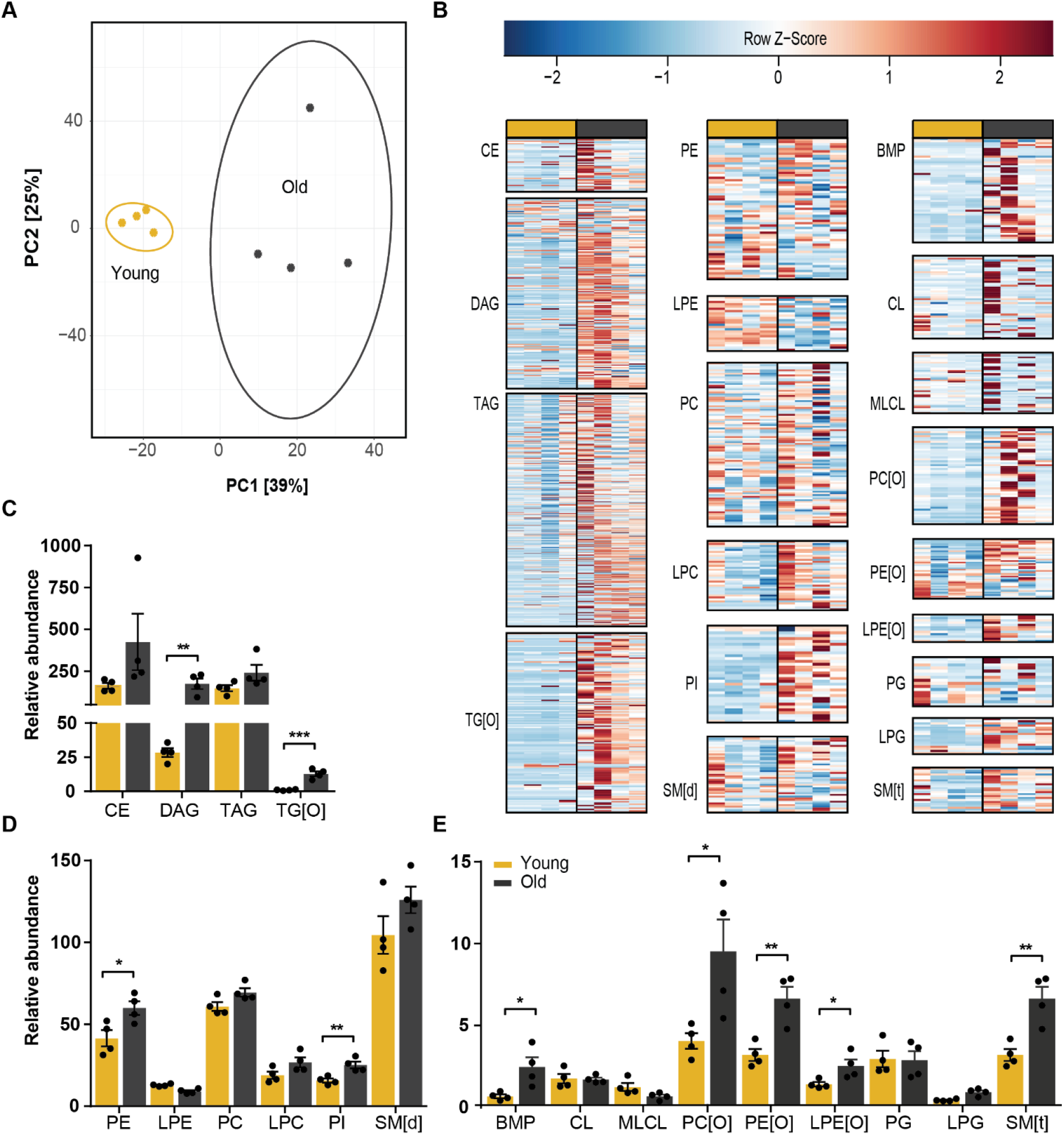
Global accumulation of lipids in aged WAT. (**A**) Principal component analysis of young and aged mice WAT lipidome shows clear separation. (**B**) Heatmaps of all lipid classes arranged from left to right by lipid abundance and ordered from top to bottom by acyl chain length. Relative abundance of (**C**) sterol and glycerolipids; (**D**), (**E**) glycerophospholipids and sphingomyelins show accumulation of almost all lipid classes. \**P* <0.05, \*\**P* <0.01, \*\*\**P* <0.001 (unpaired Student’s t-test).

Although the BAT lipidome was less affected by aging (**Figure 3A**), there were some striking differences. DAGs increased by a two-fold and BMPs were increased a four-fold in BAT from old mice (**Figure 3C-E**). Remarkably, we found that almost all BMPs, irrespective of their fatty acid composition, were higher in aged BAT (**Figure 3B**). Furthermore, TAG species tended to accumulate in aged BAT in a similar fashion as was observed in WAT. Although the total TAG levels were not significantly different, we found that TAGs composed of saturated or unsaturated fatty acids that were short (≤C14) or large (≥C24) accumulated upon aging (**Figure 3B**).

**Figure 3:**
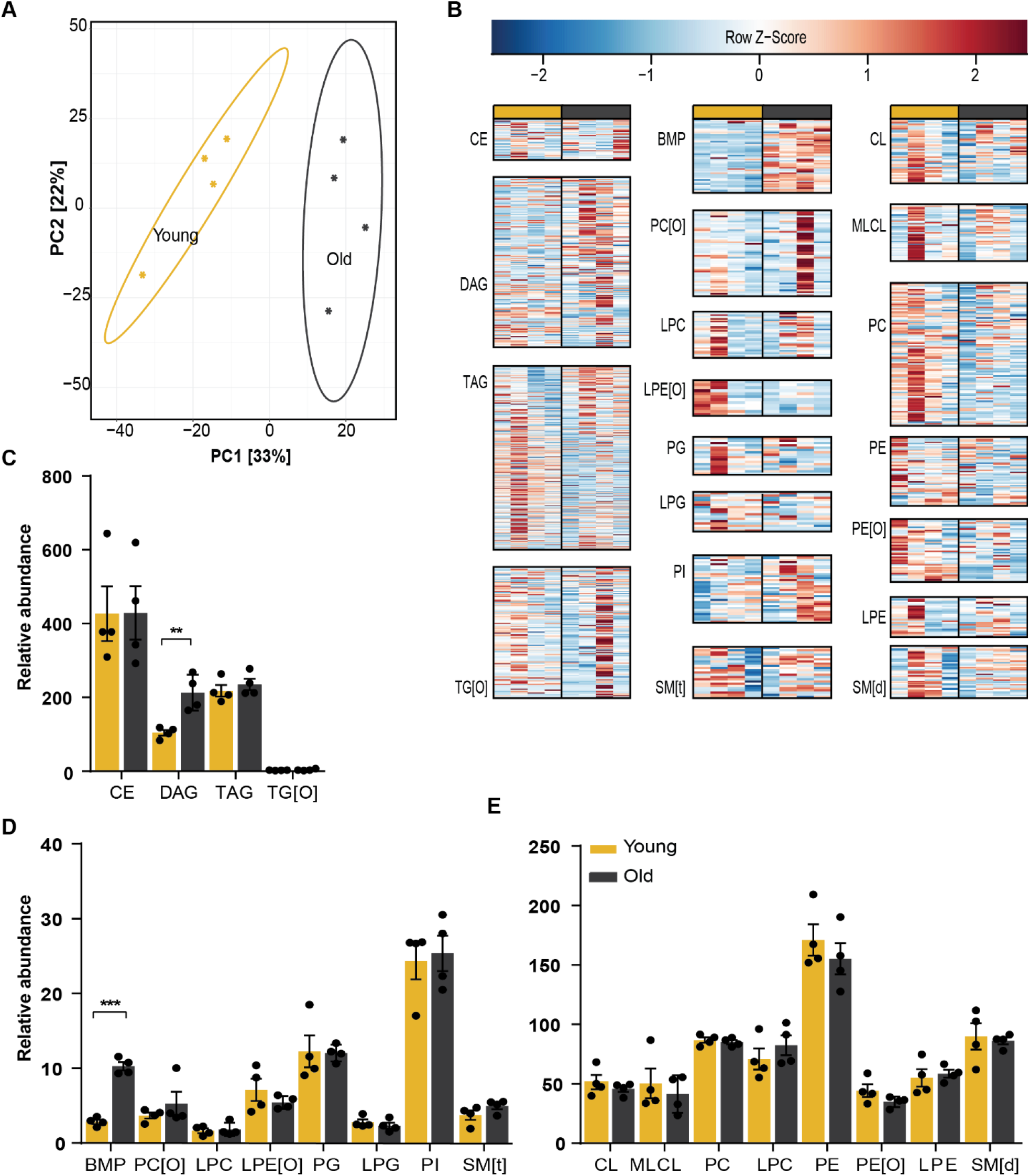
Aged BAT accumulates DAG and BMP. (**A**) Principal component analysis of young and aged mice BAT lipidome shows clear separation. (**B**) Heatmaps of all lipid classes arranged from left to right by lipid abundance and ordered from top to bottom by acyl chain length. Relative abundance of (**C**) sterol and glycerolipids; (**D**), (**E**) glycerophospholipids and sphingomyelins show accumulation of BMPs and DAGs. \*\**P* <0.01, \*\*\**P* <0.001 (unpaired Student’s t-test).

### 3.3 Lipids in aged WAT are less rhythmic than in young WAT but show rhythmic phase advancements

We next set out to investigate how aging affects the rhythmicity of the lipidome in WAT and BAT. We profiled the lipids of both adipose tissues of young and old mice at ZT2, 6, 10, 14, 16 and 22. Lipids that were above the limit of detection across all time points were included in the JTK_CYCLE analysis. In young WAT, we could detect 908 lipids of which 188 lipids were oscillating (21%; **Figure 4A**), which was reduced to 14% in old WAT (141 out of 1041; **Figure 4C**). More than half of the rhythmic lipidome of young WAT consisted of TAGs (63%), about 23% were glycerophospholipids, 4% were DAGs and 2% were sphingomyelins (SMs) (**Figure 4A**). One-third of the total TAG pool was found to be rhythmic (**Figure 4B**), suggesting that TAG metabolism is under ample temporal regulation. The percentage of rhythmic lipids from the total pool of TG[O]s, PEs, PGs and PIs was around 17%. The distribution of rhythmic lipids classes in old was different than in young WAT. TAGs remained the major group of rhythmic lipids, although reduced to 38%. The second largest group of oscillating lipids were the glycerophospholipids, accounting for around 33% of the rhythmic lipidome in aged WAT (**Figure 4C**). Finally, there was a marked increase in the number of rhythmic DAG and CE lipids (DAGs: from 4% to 8%; CEs: from 0% to 3.5%) (**Figure 4C**). For most of the lipid classes, about 8-15% of the total lipid pool was rhythmic. For PIs this percentage was doubled to about 30%, making it the most rhythmic lipid class (**Figure 4D**).

**Figure 4:**
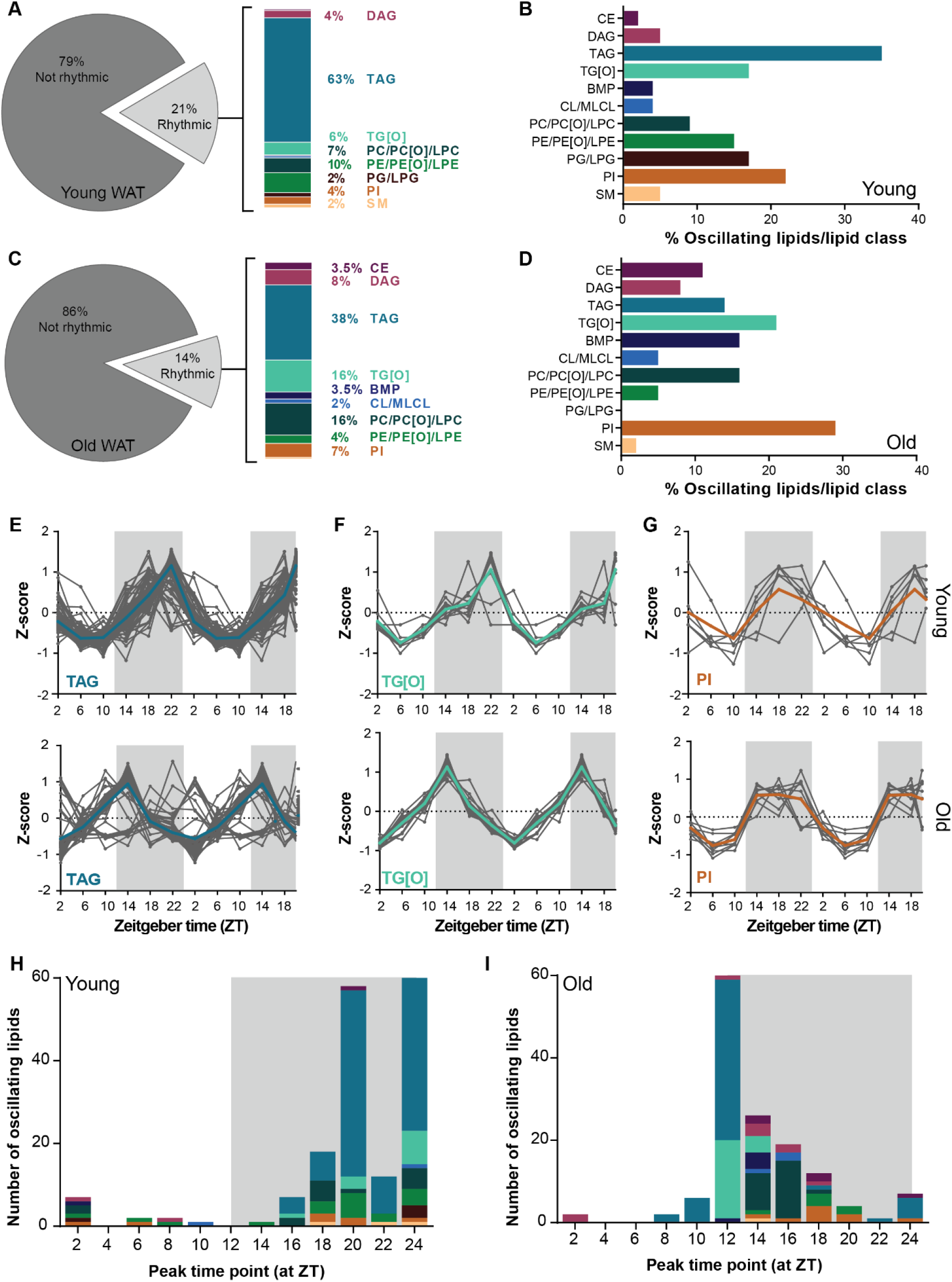
In WAT aging reduces number of rhythmic lipids and advances the lipid peak phase. (**A**) Percentage of rhythmic lipids in young and (**C**) aged mice as considered by JTK_CYCLE analysis. The rhythmic lipids were of different classes. Depiction of the rhythmic/lipid class percentage of young (**E**) and (**D**) aged mice. Double plotted rhythmic profiles of (**E**) TAGs, (**F**) TG[O]s and (**G**) PIs in young mice compared to aged mice. The gray lines represent the individual lipid species and the bold colored lines the group average. The data were double plotted to increase the visibility of day/night rhythms. The peak time point of individual lipids species depicted with the color to their corresponding lipid class of young (**H**) and (**I**) aged mice.

Aging affects SCN functioning, for instance with a reduced amplitude in electrical activity, this affects the regulation of peripheral rhythms [30]. Therefore, we next examined if aging affects the phase of rhythmic lipids in aged WAT. **Figures 4E-G** shows examples of diurnal profiles of rhythmic lipid classes (all other profiles can be found in **Supplementary Figure S1**). Side-by-side comparison of lipids in young versus aged WAT revealed that the rhythmic lipids peak at different times. For example, TAGs and their alkylated counterpart TG[O] peak early in the dark phase in old WAT, whereas the peak level in young WAT was reached at the end of the dark phase (**Figure 4H-I**). This difference was even more pronounced when visualizing at what time point each rhythmic lipid reached their peak level (**Figure 4H-I**; see also **Supplementary Table S1-S2**). In young WAT, peak levels were generally scattered during the dark phase. This was especially the case for glycerophospholipids and sphingolipids, which peak at different time points throughout the dark phase. Glycerolipids tended to peak at the end of the dark phase (**Figure 4H**). In contrast, the rhythmic glycerolipids in old WAT peaked almost exclusively at the beginning of the dark phase, with some exceptions peaking towards the middle of the dark phase (**Figure 4I**). Together, this suggest that with aging the number of rhythmic lipids in WAT are reduced and the peak phases are advanced.

### 3.4 The rhythmic lipid profile is influenced by chain length in young BAT but not in aged BAT

The number of rhythmic lipids in young BAT was larger and more diverse than in young WAT. In young BAT, we annotated 1041 lipids of which 228 were considered rhythmic (22%; **Figure 5A**). One-third of the oscillating lipids were TAGs, followed by DAGs. PGs were most rhythmic as a class, as over 40% of the total PG pool was oscillating. DAGs, BMPs and SMs also showed considerable rhythmicity with about 30% being rhythmic (**Figure 5B**). The percentage and distribution of rhythmic lipids was altered upon aging. In aged BAT, 18% of the lipidome was rhythmic which equals 202 lipids out of 1099 analyzed lipids (**Figure 5C**). A third of these were TAGs, followed by DAGs, but we also observed a striking increase in the percentage of PE and CL species in the oscillating lipids. Indeed, about 20-30% of the total PE and CL pool, respectively, was rhythmic. In general, several lipid classes that had a substantial number of rhythmic species in young BAT, were less rhythmic in old BAT. This was the case for BMP, PG, PI, and SM lipids. TG[O]s were even completely absent in the rhythmic lipid pool (**Figure 5D**).

**Figure 5:**
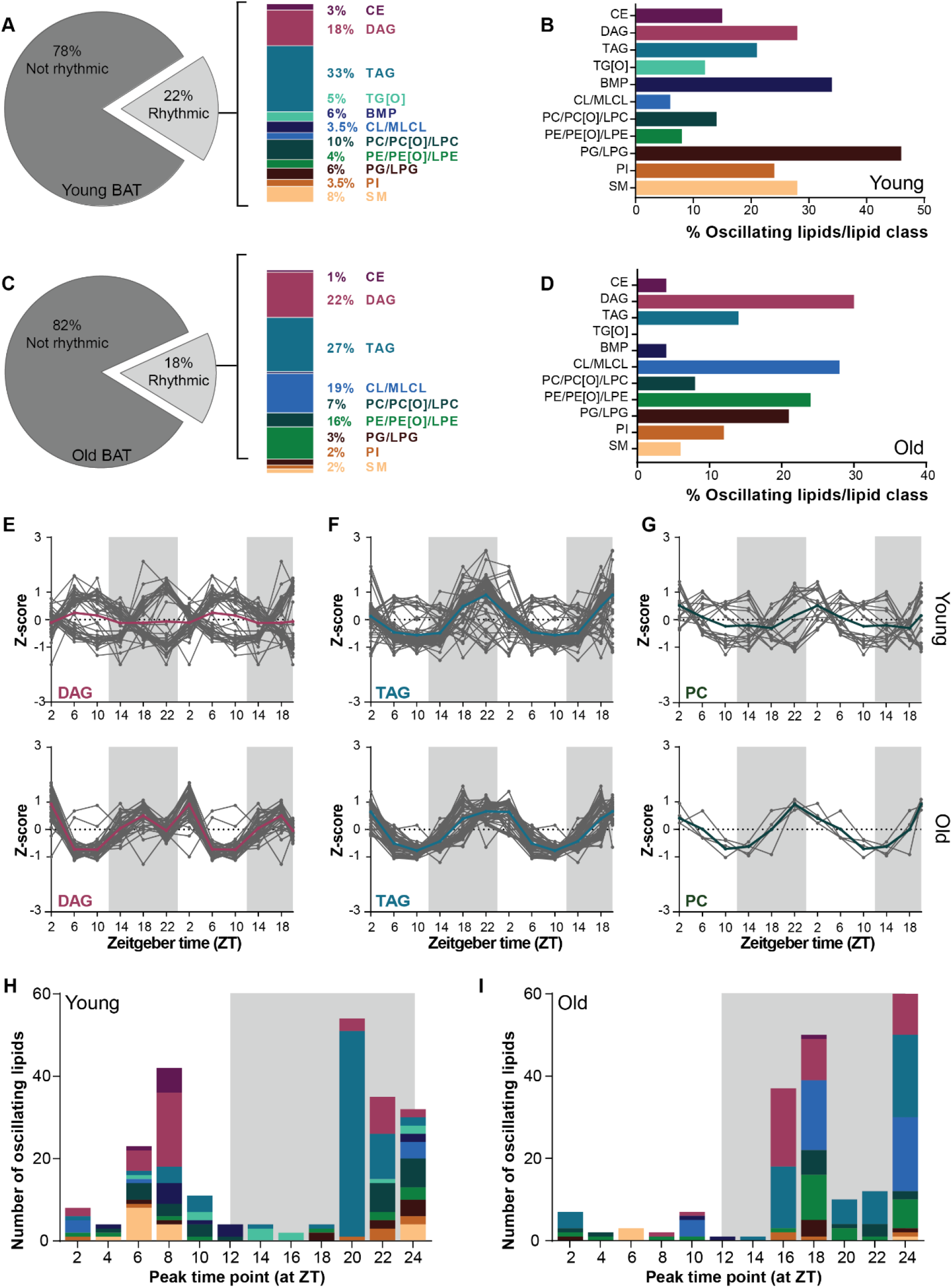
Rhythmic lipid profiles is influenced by acyl chain length in young BAT. (**A**) Percentage of rhythmic lipids in young and (**C**) aged mice as considered by JTK_CYCLE analysis. The rhythmic lipids contained lipids of different lipid classes. Depiction of the rhythmic/lipid class percentage of young (**E**) and (**D**) aged mice. Double plotted rhythmic profiles of (**E**) DAGs, (**F**) TAGs and (**G**) PCs in young mice compared to aged mice. The gray lines represent the individual lipid species and the bold colored lines the group average. The peak time point of individual lipids species depicted with the color to their corresponding lipid class of young (**H**) and (**I**) aged mice.

Another striking difference between the oscillating lipidome of young and aged BAT was the diurnal profile. In aged BAT, three major contributors to the oscillating lipidome, TAG, DAG and PC, followed opposing profiles in which some species peaked during the light phase while others peaked during the dark phase (**Figure 5E-F**). Interestingly, this difference in peak time appeared to be related to the acyl chain length of the rhythmic species these reached their peak levels. In general, rhythmic lipids that peaked around ZT6-10 contained fatty acids with acyl chain of ≥C18:x (e.g. ≥(36:x) for diarylspecies and ≥(54:x) for triarylspecies), while those containing acyl chain of <18 carbons (e.g. <(36:x) and <(54:x)) peaked at around ZT22 (**Supplementary Table S3**). Many of these larger lipid species were either not present or differentially regulated in the rhythmic lipidome of aged BAT (**Supplementary Table S4**). The majority of rhythmic lipids in old BAT reached their peak level during the dark phase, while in young BAT this was more diverse with some lipids peaking during the middle of light phase and others towards the end of the dark phase (**Figure 5H-I**).

### 3.5 Rhythmic lipidome is differentially regulated in aged adipose tissue

Periodic regulation appears to have a distinct effect on the WAT and BAT lipidome, and this regulation seems to be further modified during aging. To dissect the tissue-specific effects of aging process we cross-compared the individual lipids from the rhythmic lipidome of WAT and BAT in young and aged mice (**Figure 6A**). In WAT, only 12 out of 305 detected rhythmic lipids were rhythmic in both young and old tissue. Seven of these were unique to WAT, this included three TG[O]s, two TAGs, and a single PC and PI, which all contain fatty acids of ≥C18:x (**Figure 6B**). In BAT, there was more overall diversity among the rhythmic lipids. We found 51 lipids out of 328 that were rhythmic in young and aged mice, 26 of which were unique to BAT (**Figure 6B**). The majority of the rhythmic lipids were small DAG species (≤(32:x), composed of two fatty acids ≤C16) that peaked during the dark phase (**Supplementary Table S3-5**). Other lipids only rhythmic in BAT were glycerophospholipids of various classes. In total, we detected 416 lipids that were rhythmic either in WAT or in BAT. Little over 10% (45 lipids) of the rhythmic adipose lipidome was universal to young adipose tissue. The lipids that we found in young WAT and BAT were mostly shorter TAGs that peak at the end of the dark phase (**Figure 6B**, see also **Supplementary Table S1** and **S3**). Many of these lipids were lost upon aging. We observed overlap of only four lipids in old WAT and BAT. Finally, PI(34:2) and PI(38:2) were the two lipids that showed universal rhythmicity in all tissues at all ages.

**Figure 6:**
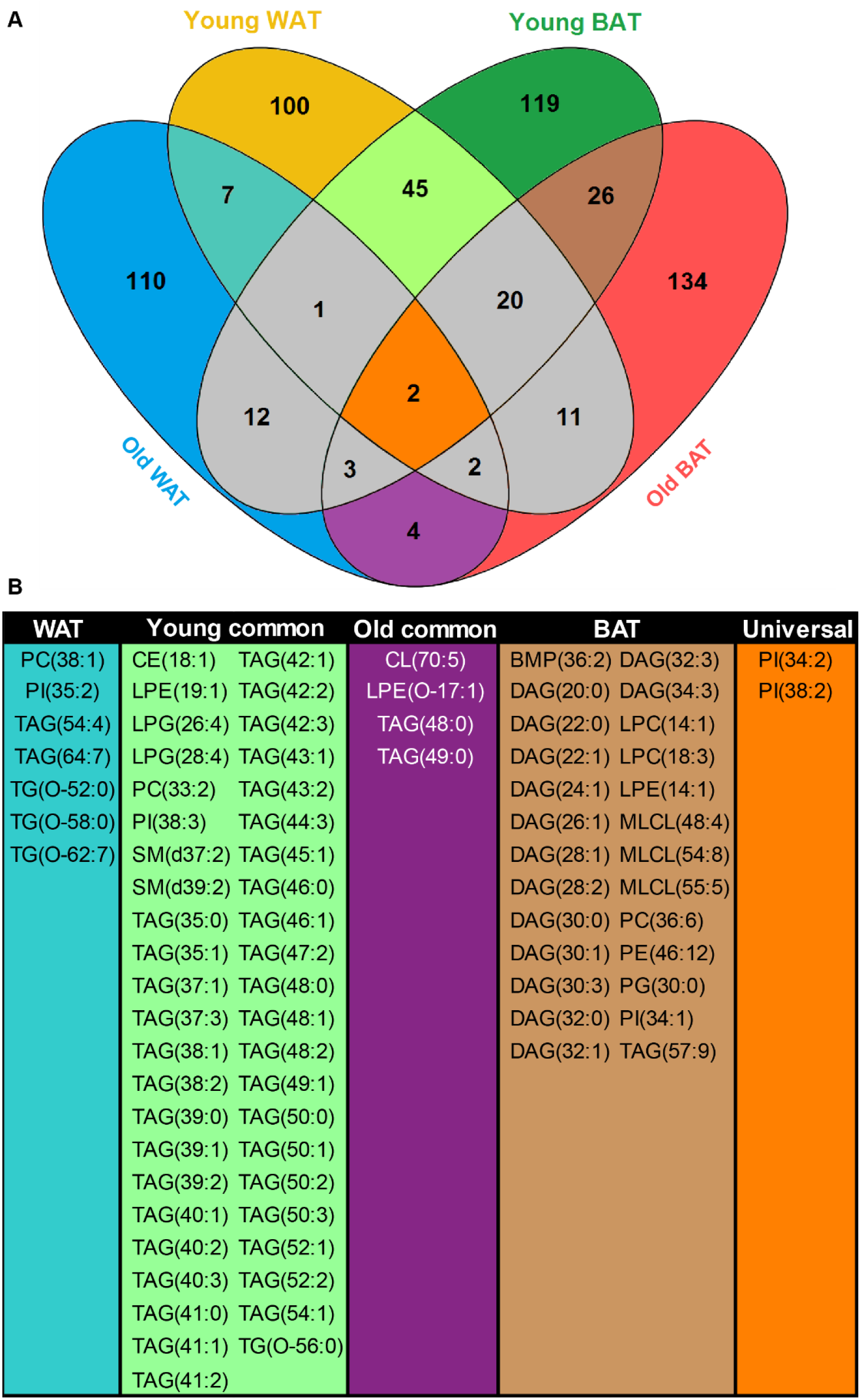
Overlap of rhythmic lipids in WAT and BAT upon aging. (**A**) Venn diagram depicts the number of unique and shared rhythmic lipid species of young and old WAT and BAT. The interesting areas of overlap and the number unique lipids are depicted in color. (**B**) Individual lipids species that are rhythmic in young and old WAT(WAT); in young WAT and BAT (Young common); in old WAT and BAT (Old common); in young and old BAT (BAT) and in all groups (Universal) as depicted in Venn diagram.

## 4. Discussion

The relation between the circadian system and aging has long been acknowledged, however the direction of this relationship remains elusive [7]. What is generally accepted, is that aging affects the entrainment of the central clock in various ways, and that dyssynchrony in the central clock can cause dysregulations in the peripheral clocks [31]. What is still under debate, however, is if peripheral clocks are individually affected upon aging, given that many age-induced effects appear to be tissue-specific [30]. In this study, we provide a global analysis of lipid species in WAT and BAT around the clock, in young and old mice. We have characterized the oscillating lipidome in WAT and BAT and show that the number of rhythmic lipids diminish upon aging.

Whole body knock out of one the central clock transcription factors, BMAL1, has been found to accelerate aging and induce metabolic disorder [5,32]. The effect of premature aging due to BMAL1 deletion was previously studied in different adipose depots, and revealed adipose deterioration and overall reduction of adiposity upon aging [33]. However, adipose specific disruption of BMAL1 caused mice to become obese [34]. The tissue-specific deletion also affected WAT and BAT lipid levels in different manners; WAT showed reduced content of poly-unsaturated fatty acids whereas in BAT the fatty acyl chain length was reduced [35]. The authors explained these phenomena by the reduced expression of *Elovl6* and *Scd1*, which are genes encoding enzymes involved in fatty acid elongation and desaturation. These results highlight that the interplay between aging and the circadian system are tissue- and context-dependent. In our study, aging was accompanied by a general increase of lipids upon aging in both WAT and BAT. One of the most striking lipid increases in both adipose tissues was that of DAGs. In adipose tissue particularly, DAGs are highly bioactive metabolites, which are at the branch-point between phospholipid and neutral lipid metabolism. For example, DAGs can be formed through lipolysis of TAGs or esterification of fatty acids to glycerol, but can also be synthesized from lipid precursor phosphatidic acid [36]. Interestingly, despite the abundance of TAGs, the average TAG level was not significantly increased in aged adipose tissue. However, in both adipose tissues the shorter chain and the longer chain TAGs tended to accumulate. In a recent study focused on lipid metabolism, a general accumulation of glycerophospholipids was also found in aged BAT especially, while TAGs declined with aging [37]. Furthermore, enzymes involved in DAG metabolism could be upregulated upon aging, which would favor DAGs to increase without affecting or even reducing TAG levels. For instance, expression of the adipose triglyceride lipase (*Atgl*; triglyceride lipolysis) but also diacylglycerol acyltransferase-1 (*Dgat1*; triglyceride esterification) increase in WAT depots upon aging [38]. Interestingly, the adipose master regulator PPARγ and its many downstream enzyme effectors—including those involved in DAG and TAG metabolism such as DGAT, ATGL, hormone sensitive lipase and phosphatidic acid phosphatase enzymes—exhibit circadian oscillations [9,39,40].

The relationship between circadian gene expression and aging was recently explored in the liver. Sato *et al*. compared the hepatic circadian transcriptome of young and old mice and described that although expression of core clock was not significantly affected, circadian expression of many genes, especially those involved in metabolism, were altered [15]. Previously, about 13% of the WAT and 15% of the BAT gene program was found to exhibit a conserved circadian expression profile and many of these genes are involved in metabolism [41]. A recent metabolomics study showed that less than 10% of the WAT metabolome and up to 50% of the BAT metabolome is rhythmic [42]. Many of these rhythmic metabolites were lipids. In the present study, we found that ~18% of the adipose lipidome is rhythmic. Rhythmicity of specific lipid classes was not always in correspondence with the increase of said lipid class. For example, in WAT a plethora of lipids accumulated (i.e. DAG, TG[O], PE, PI, BMP, PC[O], PE[O], LPE[O], and SM[t]), yet only the CE, DAG, BMP and PI lipids maintained or showed altered rhythmicity. In BAT, BMPs accumulate in old mice yet became less rhythmic. On average, the number of rhythmic lipids decreased with 2-5 percentage points upon aging, suggesting that the processes that cause age-associated lipid accumulation interfere with the regulation of diurnal lipid concentration. In contrast, the CL/MLCL concentration was not affected by aging, but the number of rhythmic CL/MLCL lipids increased in aged BAT.

Adipose tissue function is regulated by the circadian clock. During fasting adipose tissue should release fatty acids into the circulation to be used as energy substrates. Likewise, fatty acids are also required upon prolonged physical activity. On the other hand, during feeding, lipids from the diet are taken up and stored in the adipose tissue. It is therefore not surprising that most rhythmic lipids found in both WAT and BAT are TAGs and most lipids specific to BAT are DAGs, which reflects the storage function of adipose tissue. BAT is a metabolically highly active organ that takes up most lipids at the beginning of the dark phase when animals become active [29]. We observed that the rhythmicity of DAGs, TAGs, and PCs in BAT is presumably influenced by this process. In our study, there was a specific diurnal regulation of lipids depending on their acyl chain length. Lipids with acyl chain lengths longer than C18:x peaked during the light phase, while shorter chain lipids peaked during the dark phase. According to BAT lipid uptake activity, it is most likely that lipids that peak during the dark phase are lipids taken up from the diet. The longer lipids that peak during the light phase are most likely the result of fatty acid elongation followed by incorporation in lipids. This is corroborated by the expression patterns of genes coding for enzymes involved in de novo fatty acid synthesis and elongation in the liver, including *Acly, Acaca, Fasn, Elovl3, Elovl5* and *Elovl6* which are highest at the end of the dark phase [43,44]. In old mice the peak time of long-chain containing lipids changed. This is indicative of impaired diurnal regulation of lipid elongation. In general, rhythmicity of long-chain lipids was reduced in old mice. Finally, a group of glycerophospholipids that was abundant in all adipose tissue independent of age, were the PI lipids. We found two unique PI species to be rhythmic in all experimental groups. In general ~20% of the PI pool was rhythmic in both BAT and WAT, which is similar to the percentages of PI lipids found to be rhythmic in in muscle and liver [45,46]. This suggests that the diurnal regulation of PI lipids is important for adipose function. Like many glycerophospholipids, PIs are important lipids for cell membrane function. More specifically, PI lipids can play important roles in regulating metabolism, as they are signaling molecule precursors required for the PI3K pathway, which is involved in the regulation of many metabolic pathways including insulin signaling [47]. In liver, PI lipid levels were heavily affected by feeding/fasting time, which corresponds with its putative function in metabolic regulation [46]. In future research, we should further explore this relationship between diet and enzymatic regulation and their effect on adipose tissue lipid levels.

In conclusion, our study demonstrates the effects of aging on adipose tissue metabolism and its intricate relation with the circadian system. Our work contributes to the notion that intrinsic regulation and external factors such as behavior have a big effect on the circadian control of metabolism. This insight may lead to future methods of intervention to benefit circadian regulation of metabolism to improve longevity and healthspan.

## Acknowledgements

This work is financially supported by Rembrandt Institute for Cardiovascular Science (to RHH) and The Netherlands Cardiovascular Research Initiative: an initiative with support of the Dutch Heart Foundation (CVON2014-02 ENERGISE). SK is funded by the Dutch Heart Foundation (2017T016), GEJ is supported by a VENI grant from ZonMw (no. 09150161810014). RHH is funded by an ERC Starting grant (no. 638290) and a ZonMw VIDI grant (no. 917.15.305).

## Author contributions

NMH, MvW and RHH: designed the project; NMH, RB, HLE, SK: performed the experiments and analyzed the data; SM and GEJ: contributed reagents/materials/analysis tools; NMH, GEJ, FMV, RHH and MvW: interpreted and discussed the results; NMH, FMV, RHH, MvW: wrote and revised the manuscript. All authors reviewed and approved the manuscript.

## Conflict of interest

The authors report no conflict of interest related to this work.

## Supplementary material

**Supplementary Figure S1.**
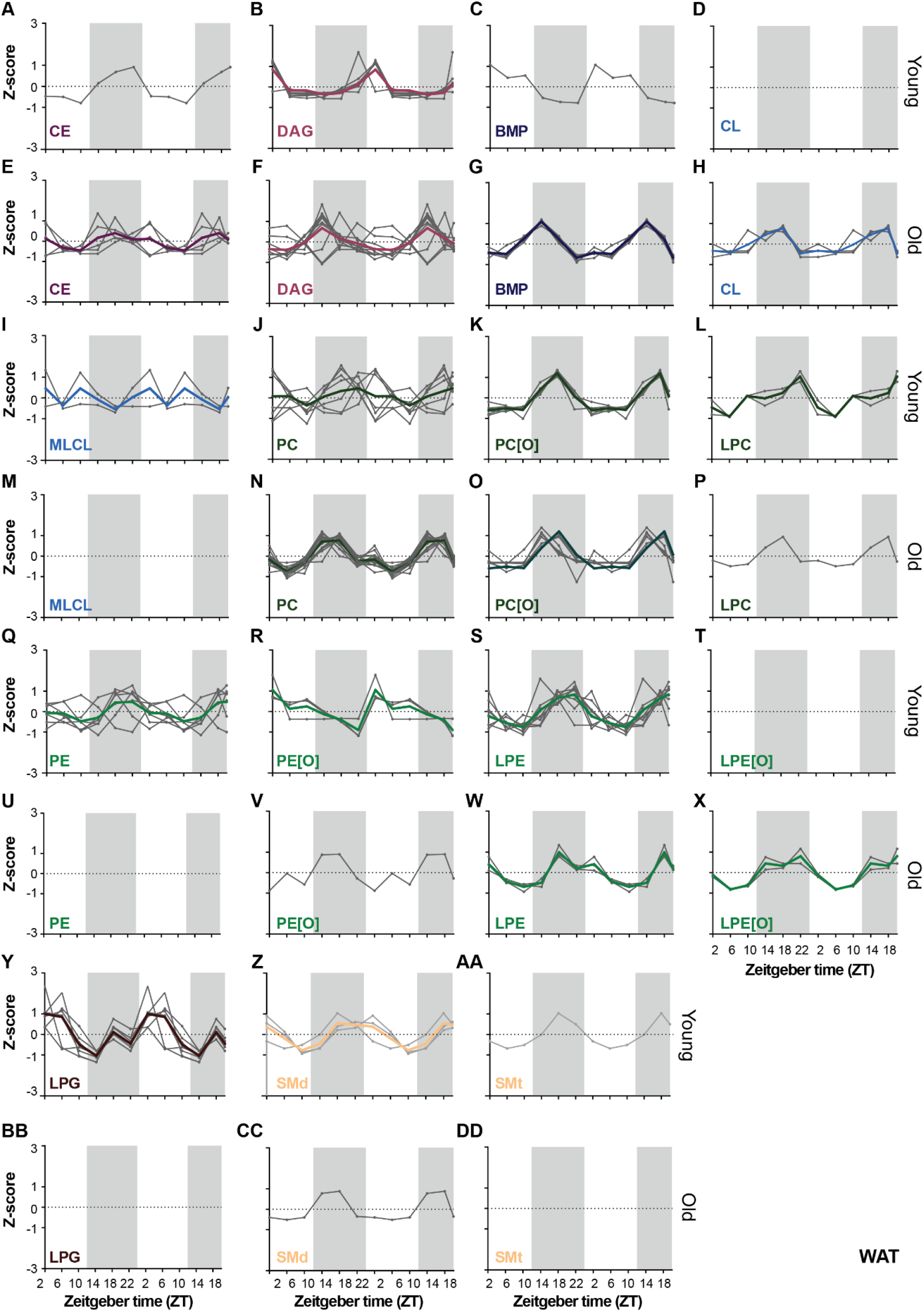
Related to Figure 4: Rhythmic lipid profiles in WAT. (**A-D**), (**I-L**), (**Q-T**), (**Y-AA**) lipid profiles of young and (**E-H**), (**M-P**), (**U-X**), (**BB-DD**) old mice. The gray lines are the individual rhythmic lipid species, the bold and colored line represents the average of the rhythmic lipid class.

**Supplementary Figure S2.**
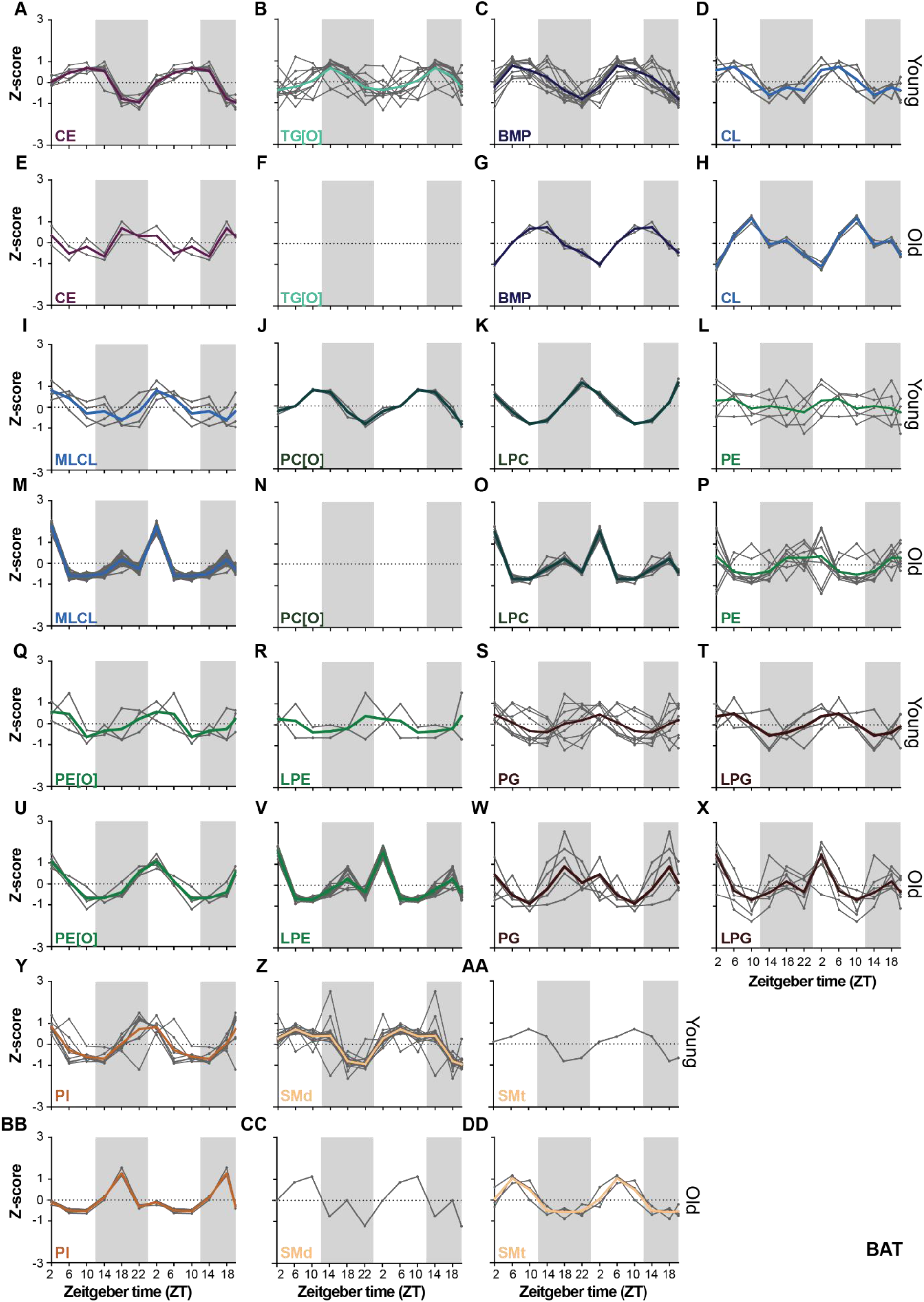
Related to Figure 5: Rhythmic lipid profiles in BAT. (**A-D**), (**I-L**), (**Q-T**), (**Y-AA**) lipid profiles of young and (**E-H**), (**M-P**), (**U-X**), (**BB-DD**) old mice. The gray lines are the individual rhythmic lipid species, the bold and colored line represent the average of the rhythmic lipid class.

**Supplementary Table S1 Related to Figure 4** Zenith time point of individual lipids in young WAT

**Supplementary Table S2 Related to Figure 4** Zenith time point of individual lipids in aged WAT

**Supplementary Table S3 Related to Figure 4** Zenith time point of individual lipids in young BAT

**Supplementary Table S4 Related to Figure 4** Zenith time point of individual lipids in aged BAT

## Notes

### Competing Interest Statement

The authors have declared no competing interest.

